# Emergent Homeostasis and Degeneracy from multi-Dimensional Attractors

**DOI:** 10.1101/2025.05.07.652484

**Authors:** Hanna Salman, Kuheli Biswas, Naama Brenner

## Abstract

Biological systems maintain homeostasis, ensuring stability in the face of internal and external perturbations and counteracting stochastic noise. Traditionally, this is understood through the lens of control mechanisms designed to offset deviations and maintain certain quantities near functionally desired set-points. Here, we propose an alternative perspective to understand homeostasis: the *dynamic perspective*, in which homeostasis emerges from high-dimensional interactions creating stable attractors in the phase space. These multi-dimensional attractor manifolds can constrain all components collectively, eliminating the need for explicit control of individual variables. The presence of null directions on an attractor allows for degenerate states that can add flexibility while preserving functionality. Using single cell growth and division homeostasis as a test case, we develop and support our perspective by models and meta-analysis of numerous single-cell datasets across organisms and conditions. Importantly, we do not reject the control-theory perspective but rather suggest that control circuit models can be seen as low-dimensional projections of a more complex, multi-dimensional system.

## Introduction

Living systems maintain stability in many of their properties despite internal and external perturbations, a phenomenon known as homeostasis [1, 2, 3]. This is a property of all scales of biological organization, from molecular circuits and cellular properties, to organs and physiological function. When homeostasis fails, system dysfunction occurs, making it a subject of significant interest and extensive research over many years. The most widely held approach to study homeostasis entails that control mechanisms have developed along evolution which offset molecular and/or environmental perturbations, and keep certain quantities near functionally desired set-point values. Control theory, with its mathematical and conceptual tradition, provides a natural framework for describing such processes, and has been utilized extensively for understanding biological homeostasis [4, 5, 6]. One drawback of this approach is that complex biological systems are typically high-dimensional, and include countless interactions among their components. In such a system, designing or even conceptualizing a set of controls that coherently regulate many specific target values and compensate for deviations in each component, is highly challenging.

However, such systems can naturally exhibit **attractors**, that collectively constrain the dynamics of multiple variables. This may obviate the need for dedicated feedback control circuits that offset perturbations in individual variables from their specific set-points. The concept of attractor generalizes that of a fixed point (a zero-dimensional attractor) to higher-dimensional structures, such as stable limit cycles or line attractors. In this opinion paper we put forward the **dynamic perspective** for understanding biological homeostasis, where stability and homeostasis can arise in a self-organized manner through intrinsic dynamics. Dynamical systems theory posits that a system’s behavior is largely governed by its phase-space topology — specifically, the nature of attractors and their stability properties. Therefore, we focus our perspective of biological homeostasis on understanding the emergence and properties of such attractors.

Collective attractors may be characterized by degeneracy, and more generally by highly non-uniform stability in different directions. For instance, if the attractor defines a geometric structure such as a smooth manifold, trajectories may evolve along directions within the manifold while remaining restricted in directions perpendicular to it. More generally, in the high-dimensional phase space, some directions may be immensely more stable than others [7, 8]. Biologically, this would imply that not all system variables are equally regulated, and different levels of homeostasis are induced on individual components. In some cases, it may imply that the system’s stable functionality is entirely degenerate with respect to some variables, suggesting that they need not be regulated at all. Interestingly, such degeneracy has been independently observed in various biological contexts [9]; we argue below that it can even contribute to homeostasis.

We focus on the biological cell, the fundamental unit of life, as a prototype system to present and develop our perspective. We explore homeostasis and variability in cellular growth and division, demonstrating how a dynamic attractor, emerging from complex cellular interactions, enforces multi-variable homeostasis. We characterize the geometric properties of this attractor, linking them to biological function, and identify two distinct forms of variability related to its stable and degenerate directions: *temporal fluctuations within a single cell*, which may destabilize the system, and *persistent variability among individuals*, which may enhance adaptability while maintaining functionality and homeostasis. The dynamic perspective and its predictions, as well as the distinction of the two types of variability, are supported by a broad set of single-cell measurements that align closely with several clear theoretical predictions. Finally, we discuss the generality of our findings and their connection to other biological systems.

### Dynamic attractors of growth and division

Recent advances in single-cell experimental techniques, allow the quantitative study of cellular growth dynamics with high statistical power and time span [10]. Single cells display significant variability in all measurable properties such as shape and size, molecular content, growth rate, etc. This variability is observed even when their genomes are identical, and they are all under homogeneous environmental conditions. At the same time, the dynamics of proliferating cell populations maintain homeostasis of growth and division over multiple cycles. Moreover, this homeostasis entails balance between multiple cellular components. These properties have highlighted several questions: what mechanisms control growth and division homeostasis? How are individual cellular components coordinated and stabilized?

Here, we explore *stable dynamic attractors* in models of cellular growth and division, investigating how they regulate temporal fluctuations and examining their underlying geometric structure. To this end, we focus on processes within a single cell, analyzing growth and division through the lens of dynamic attractors. As we will see, the dimensionality of the model plays a crucial role in determining the range of possible behaviors.

*Fixed points in one-dimensional models*. At a coarse-grained level, cell growth and division can be described using a discrete mapping from one cell cycle to the next. Recent studies often model this process by focusing on a single variable, such as cell size or protein content. Consider a cell starting with an initial size *V*_*n*_ at cycle *n*. If it multiplies by a factor *m* over the cell cycle and divides by a factor *f* at the end of the cycle, the discrete map is given by:

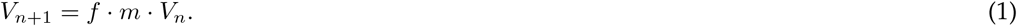

As an example, many bacteria divide symmetrically and, on average, double their size over the cell cycle, with *m* = 2, *f* = 1*/*2. Theoretically, this mapping is sensitive to fluctuations in *m* and *f* : due to the multiplicative nature of the map, such fluctuations accumulate over time, leading to instability. In the absence of other negative correlations, even small fluctuations in these parameters would accumulate and disrupt homeostasis. Hence, in this one-dimensional framework a control mechanism for cell division based on the doubling of its size is unstable.

In contrast, if we assume that the cell adds a fixed quantity of biomass or protein Δ and then divides in half, this process has a stable fixed point, allowing homeostasis to be maintained even in the presence of significant noise in Δ and *f*. This is the well-studied “adder model” [11, 12], where the mapping is described by:

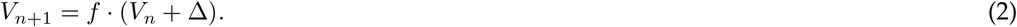

It is easy to show that, with symmetric division *f* = 1*/*2, this mapping has a stable fixed point *V* ^*\**^ = Δ. In the presence of noise both in *f* and in Δ, *V*_*n*_ converges to a stable distribution around an average size ⟨*V* ⟩ = Δ, whose properties have been studied theoretically [13, 11]. This is an example of a coarse-grained model of cell size control, where homeostasis is induced by implicit feedback arising from the attractor of the mapping. If cell division is triggered by the accumulating of some protein that adds a fixed value over the cycle, and assuming further a strong correlation between cell size and proteins - namely, balanced biosynthesis - one may motivate the adder model by known biological mechanisms [14].

This one-dimensional level of description leaves several directions for refinement. First, continuous dynamics within the cell cycle are neglected; the mapping (2) could equally reflect exponential, linear, or any other form of accumulation between divisions. We argue below that the approximately exponential accumulation observed in many cases has significant implications to the underlying dynamics.

Second, the main empirical support for the adder control model is the independence of added size on initial size, across the range of initial sizes observed experimentally. However, while Δ was found empirically uncorrelated with initial cell size - *ρ*(*V*_*n*_, Δ_*n*_) ≈ 0 on average over large data sets, a more detailed analysis that examines individual lineages separately reveals a small negative correlation over time [15]. Additionally, compensation between sister cells was observed by tracking them for a full cell cycle after separation; the smaller sister exhibits, on average, a larger added volume [16, 17, 18], again in contradiction to the principle of fixed added size as a control trigger for division.

Finally, the exponential accumulation mentioned above characterizes not only cell size but also highly expressed proteins. The rate of this accumulation fluctuates across cycles [15, 19, 20, 21, 12, 22, 23], with strong correlations observed among various cellular components [15, 20, 23]. This correlation provides a more detailed and quantitative characterization of balanced biosynthesis, or multi-variable homeostasis, which calls for a deeper understanding [15].

But how does such balance emerge among the many proteins and other cellular components? Is it the result of precise co-regulation, or could it arise as an example of emergent stability induced by complex interactions that give rise to a collective dynamic attractor, as discussed earlier? To address this, we now examine high-dimensional, continuous-time models of cellular growth and division, focusing on the conditions for emergence of dynamic attractors.

*Attractors in high-dimensional models*. Balanced exponential growth can naturally emerge in a network of coupled chemical kinetics, as observed many years ago [24]. An arbitrary system of coupled linear reactions typically converges to exponential growth of all components at the same rate. This occurs because, in linear systems, the largest eigenvalue of the interaction matrix dominates as time progresses; this makes up a dynamic attractor - as time goes by, all trajectories are attracted towards a manifold of exponential growth at the same rate. In systems with many variables and randomly chosen interactions, the presence of at least one positive eigenvalue is highly likely, making exponential growth the typical long-term behavior [15]. While this growth is universal, the specific rate *λ* (inverse of the largest eigenvalue) depends on the details of the interactions and the parameter values.

Recent theoretical work demonstrates that dynamic attractors of balanced exponential biosynthesis emerge also in a broader class of models with *non-linearly* interacting components. This result, observed in several specific models [25, 23, 26], was found to be robust with respect to details of the interactions and the number of variables, and characterizes a large class of kinetic reaction networks under quite general conditions. Specifically, any system of interactions 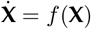 obeying the scalability condition:

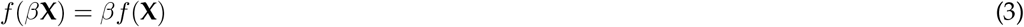

belongs to this class. If the total system volume is a linear combination of components *V* = Σ*ρ*_*i*_*X*_*i*_, all mass-action kinetic interactions are included (as can be seen by formulating them in terms of concentrations rather than quantities. For further details, see [27, 26]). Thus, similar to the linear case, a nonlinear network of chemical reactions in a growing volume will typically exhibit long-term dynamics where all components accumulate exponentially with the same rate, i.e. in balanced exponential biosynthesis. The generality of this result presents a robust theoretical basis, which motivates the use of this class of models to understand cellular dynamics in a general framework.

In the balanced state, the ratios between any two components growing at rate *λ*, remain fixed along the attractor, ensuring coordinated growth: if *X*_1_ and *X*_2_ are two such components,

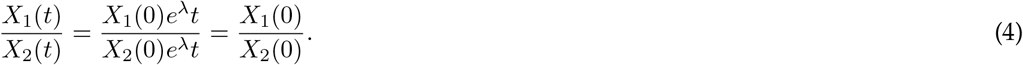

Cell size - a combination of all its components - also increases exponentially with the same common rate. As a result, molecular concentrations (i.e. the copy number of each component per unit volume) remain fixed on the attractor. Fig. 1(a) shows the exponential attractor of a specific model system [27, 23, 28], as a black solid line in the space including two out of the three dynamic variables and time. The exponential growth of each component is depicted as grey solid lines on the projection planes of the component and time, (*X*_1_, *t*) and (*X*_2_, *t*). The projection on the two components (*X*_1_, *X*_2_) is a straight line representing the fixed ratio between *X*_1_ and *X*_2_ on the attractor.

**Figure 1.**
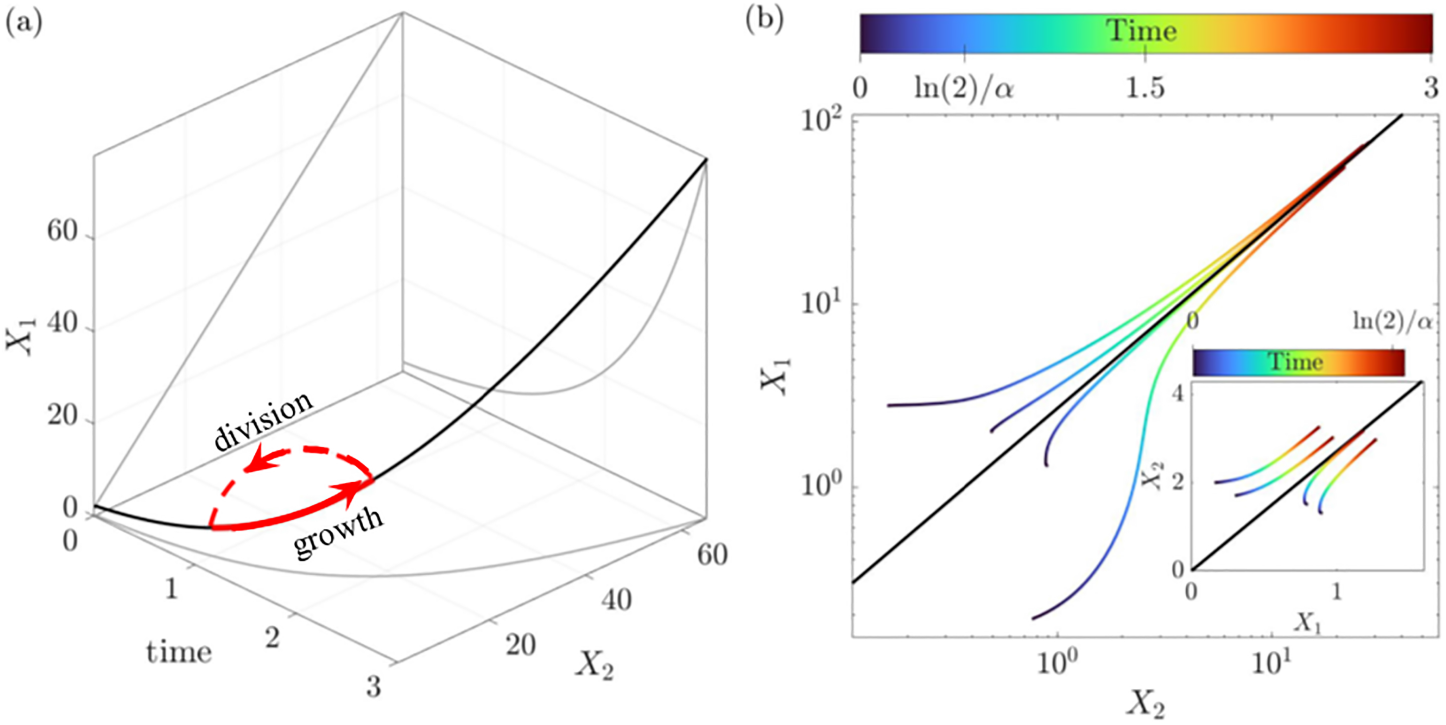
Exponential attractor in high-dimensional phase space of cellular dynamics. **(a)** Black line: The exponential attractor of the system. The projections on (*X*_1_, *t*) and (*X*_2_, *t*) planes demonstrate the exponential growth; the projection on the (*X*_1_, *X*_2_) plane highlights a property of this attractor: any two components maintain a fixed ratio along time. **(b)** Starting from arbitrary initial conditions, dynamic trajectories are drawn closer to the exponential attractor as time progresses. A typical time of the exponential growth, the doubling time ln(2)*/α*, is marked on the time (color coded); note the logarithmic axes, and the long timescale for convergence. Inset: Trajectories with arbitrary initial conditions drawn in linear scale and over the time interval 0 to ln(2)*/α*. The figure shows simulations from a specific model; details in [28].

Unlike a fixed point, the attractor is inherently dynamic, with time evolution proceeding along it at an exponential rate *λ*. This time evolution is unstable in the sense that all variables diverge to infinity at long times, but the attractor itself is a stable structure in phase space, continuously drawing nearby trajectories toward it; in this sense it is a generalization of a fixed point. The stability of the attractor is demonstrated by examining trajectories originating from various arbitrary initial conditions. Figure 1(b) illustrates these trajectories in the (*X*_1_, *X*_2_) plane (color coded for time), with the asymptotic attractor depicted by a black line. As time advances, the trajectories converge increasingly closer to the attractor. Note that the trajectories are depicted in logarithmic scales; the doubling time, a characteristic timescale of the dynamics, is marked in the color-coded representation. The inset of Fig. 1(b) presents the same trajectories on a linear scale, terminated at the doubling time.

Applying this theoretical insight to a biological cell, we note that continuous growth proceeds for a limited time—approximately doubling its constituents—before division occurs. For perfect division, all components are distributed in proportionality, thus preserving ratios among them and keeping the system’s trajectory along the attractor. This results in a hybrid continuous-discrete cycle of growth and division, in which a cell can, in principle, persist indefinitely. This cycle is illustrated in Fig. 1(a) in red. During each cycle, continuous, balanced exponential growth occurs along a finite segment (solid red line) of the theoretically infinite attractor (solid black line). At the discrete event of division, the system “jumps” back to its initial state (dashed red line), ready to begin the next cycle. Throughout this process, all components remain finite over time, and all ratios are fixed.

In reality, the growth and division of a single cell or lineage exhibit stochastic behavior over time. Division is not precise, the instantaneous exponential growth rate fluctuates, and the effective growth rate over an entire cell cycle varies across cycles. We next discuss stochastic effects on the exponential attractor and in particular on the finite-time hybrid growth and division cycle. We specifically identify two distinct types of variability, as described next.

### Type 1 variability: temporal fluctuations restrained by the dynamic attractor

Temporal fluctuations arise from many different sources, and can cause a perturbation to the perfect growth-division cycle on the attractor. Hhe stability of the dynamic attractor ensures that, following deviations, trajectories are pulled back toward it. Mathematically, this stability is reflected in the presence of negative eigenvalues of the Jacobian matrix around the attractor [15, 27, 26, 23, 28]. The existence of a stable attractor enables homeostasis across many variables simultaneously, mitigating temporal fluctuations without the need for their direct estimation or compensation of components individually, and without the need for a dedicated coordination mechanism between them. Evidence of this emergent homeostasis can be identified by analyzing the statistical properties of experimental single-cell data. Below, we present three key predictions regarding these statistical properties and demonstrate their validation through experimental data. This part summarizes and extends the results published in [28].

The first prediction arises from deviations from perfectly symmetric cell division, which can distribute various cellular components differently among daughter cells. This stochastic nature of division randomly resets the starting points of cell cycles to different nearby locations in phase space. As a result, the smooth growth along the fixed-ratio attractor is disrupted, temporarily perturbing the concerted evolution of cellular variables and causing deviations from the attractor. This deviation leads to fluctuations in instantaneous growth rates at the beginning of each cycle. However, as the cell cycle progresses, trajectories gradually return to the attractor, and growth rates stabilize, converging nearer to the steady-state growth rate characterizing the attractor. This would imply a decreasing variability in instantaneous growth rate across individual cell cycles.

To test this prediction, we estimate the coefficient of variation (*CV*_*α*_)—the ratio of the standard deviation to the mean—of the instantaneous growth rate across cell cycles as a function of cycle progression. Figure 2(a) presents the results for three different cell types, including both microbial and mammalian cells. The observed decreasing CV of the growth rate throughout the cycle provides strong experimental validation of our theoretical expectation.

**Figure 2.**
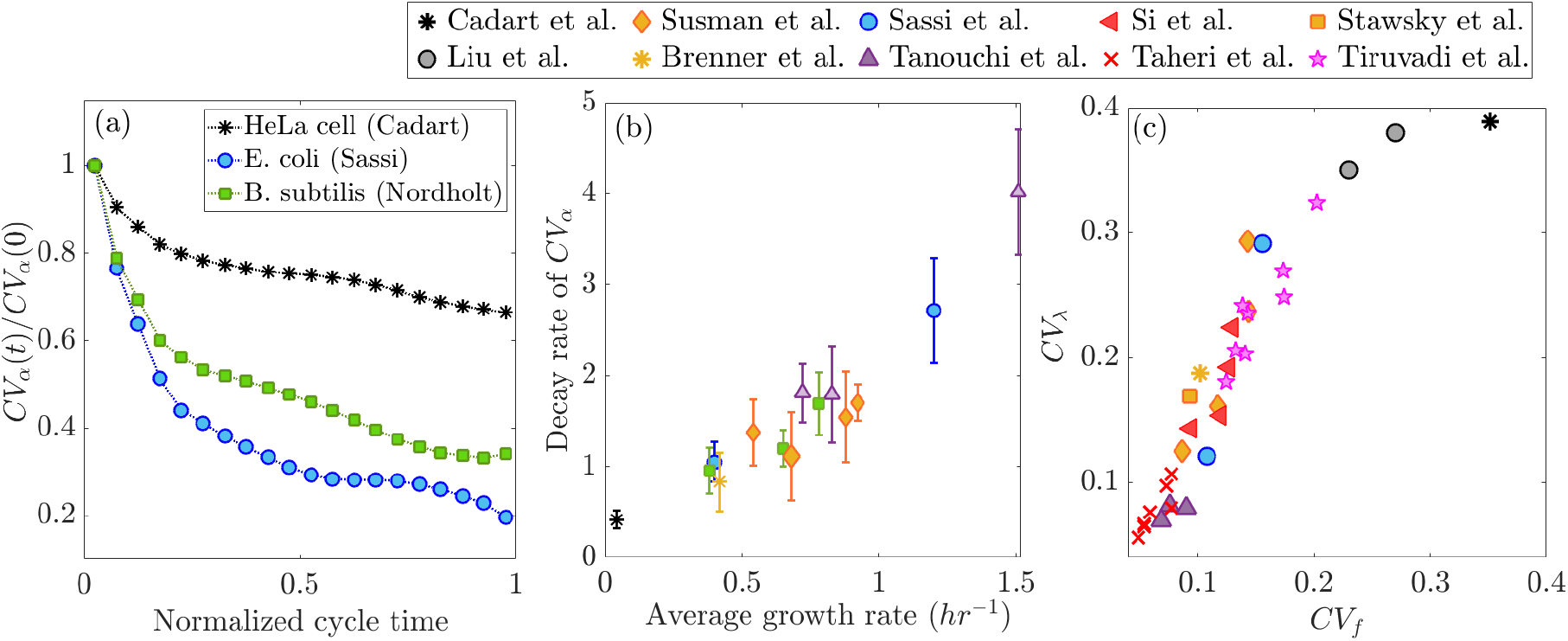
Experimental data support predictions of dynamic perspective on statistical properties of growth rate. (**a**) Decrease of CV (standard deviation over mean) in growth rate, over normalized cycle time, reflects the random initial conditions following noisy division, and the attraction of trajectories over time towards the exponential attractor. Data from *E. coli* [23] and *B. subtilis* [21] bacteria and mammalian cells [29]. **(b)** The decay rate of *CV*_*α*_ is derived from an exponential fit to the curve in (a), which increases with the average growth rate. Each point represents one experiment that includes multiple lineages; the symbols are the average and the error bars represent the standard deviation over lineages. **(c)** According to the dynamic picture, a noisy division is a primary source of disruption to the exponential attractor and thus a major driver of growth rate variability. The data were collected from various experiments, including bacterial and mammalian cells grown under diverse environmental conditions [15, 30, 20, 12, 14, 22, 23, 19, 21, 29, 31].

The second prediction, derived from the dynamic perspective, concerns the rate of return to the attractor following division, quantified by the rate of decrease of *CV*_*α*_ in Figure 2(a). In specific theoretical models, this rate was computed and found to increase with the attractor growth rate [28]. This result is intuitive, since faster growth leads to quicker progression along trajectories, thereby accelerating the return to the attractor.

Experimental validation of this prediction is presented in Figure 2(b). The figure shows the decay rate, obtained by fitting an exponentially decreasing function to the trends observed in Figure 2(a), plotted as a function of the average growth rate. Each point represents an independent experiment encompassing multiple cell lineages; the symbols indicate the mean, while the error bars denote the standard deviation across lineages. These data are drawn from seven different experimental groups and span a diverse range of single-cell measurements, including bacterial and mammalian cells grown under various environmental conditions. They reveal a clear increasing relationship between the average growth rate and the rate of decrease in variability, supporting the second theoretical prediction.

The third prediction directly relates growth rate variability to division noise. Here, we examine the effective growth rate over the cell cycle, which is determined by fitting the accumulation dynamics over the entire cycle with an exponential function. In our dynamic framework, division noise perturbs the evolution of cellular components, causing their trajectories to deviate from the precise exponential growth attractor. Nonetheless, since the dynamic range over one cycle is relatively small (approximately a factor of two), a simple exponential function with an effective exponent still provides a good approximation to the measured time trace. Its fitted value, however, will fluctuate depending on the magnitude of the deviation from the attractor at division: larger division noise is expected to cause stronger disruptions to evolution along the attractor, leading to greater deviations in the trajectory and, consequently, larger variability in the effective growth rate. Indeed, Figure 2(c) confirms this relationship across a broad range of experimental data, spanning multiple organisms and cell types, and covering a large dynamic range in both axes. Each point represents an individual experiment, where the *CV* for both division noise (*CV*_*f*_) and growth rate variability (*CV*_*α*_) were computed across the entire ensemble of cell cycles.

The above analysis suggests that cell division is a major disruption to the dynamic attractor. This leads to the conclusion that preventing cell division should extend the relaxation to the attractor, allowing the trajectory to reach closer to the attractor and thereby reducing growth rate noise further as time passes. In that case, filamentous bacterial cells that elongate without cytokinesis due to inhibited division provide another prediction: these cells should exhibit less growth rate variability than normally dividing cells as they continue to grow without division. This is consistent with recent findings in [32, 28], which align well with our theoretical framework.

These validated predictions reinforce the concept of a multi-dimensional dynamic attractor characterized by a fixed growth rate, concentrations and ratios. In this framework, homeostasis of all cellular components emerges naturally from the attractor’s stability to perturbations over time. Next, we explore additional properties of the exponential attractor, particularly its role in persistent variability.

### Type 2 variability: the degenerate structure of the exponential attractor

Careful examination of single-cell lineage growth and division data reveals the presence of not only temporal fluctuations but also persistent variability among individual cells [15, 30]. This second type of variability, which we refer to as “type 2” or persistent variability, is often overlooked. It becomes evident when comparing lineage averages over time, which reveals that some of these averages can be very different among lineages. Growth rate and cell size, in particular, are highly sensitive and display significant variation in their time-averaged value across different lineages tracked within the same experiment and even in the same trap [30]. Notably, this persistent variability does not accumulate and thus does not disrupt homeostasis; instead, each lineage can maintain growth and division homeostasis with distinct average for many generations. We argue that this inherent type of degeneracy is naturally explained by the properties of the dynamic attractor.

The exponential attractor, combined with multiplicative cell division, introduces an intrinsic degeneracy in balanced biosynthesis with respect to absolute quantities. In Fig. 1(a), one growth and division cycle is depicted (red lines), but others can arise from different initial conditions along the attractor. Thus, while perpendicular deviations are constrained by the attractor stability, a flat direction permits lineage-specific variation; lineages can coexist around different segments of the attractor resulting in distinct time-averaged absolute quantities such as cell size and protein content.

To illustrate this effect, we simulated two lineages following the model dynamics, starting from different initial conditions and incorporating noisy division. In Fig. 3(a), the exponential attractor is depicted by a heavy line, marking a fixed ratio between the two components (see projections in Fig. 1. The two lineages, (thin colored lines) each remain close to the attractor across for multiple cycles, but occupy distinct regions. As discussed earlier, due to the limited dynamic range covered within a doubling time, individual cycle trajectories do not fully return to the attractor but are still influenced by it, remaining in its vicinity.

**Figure 3.**
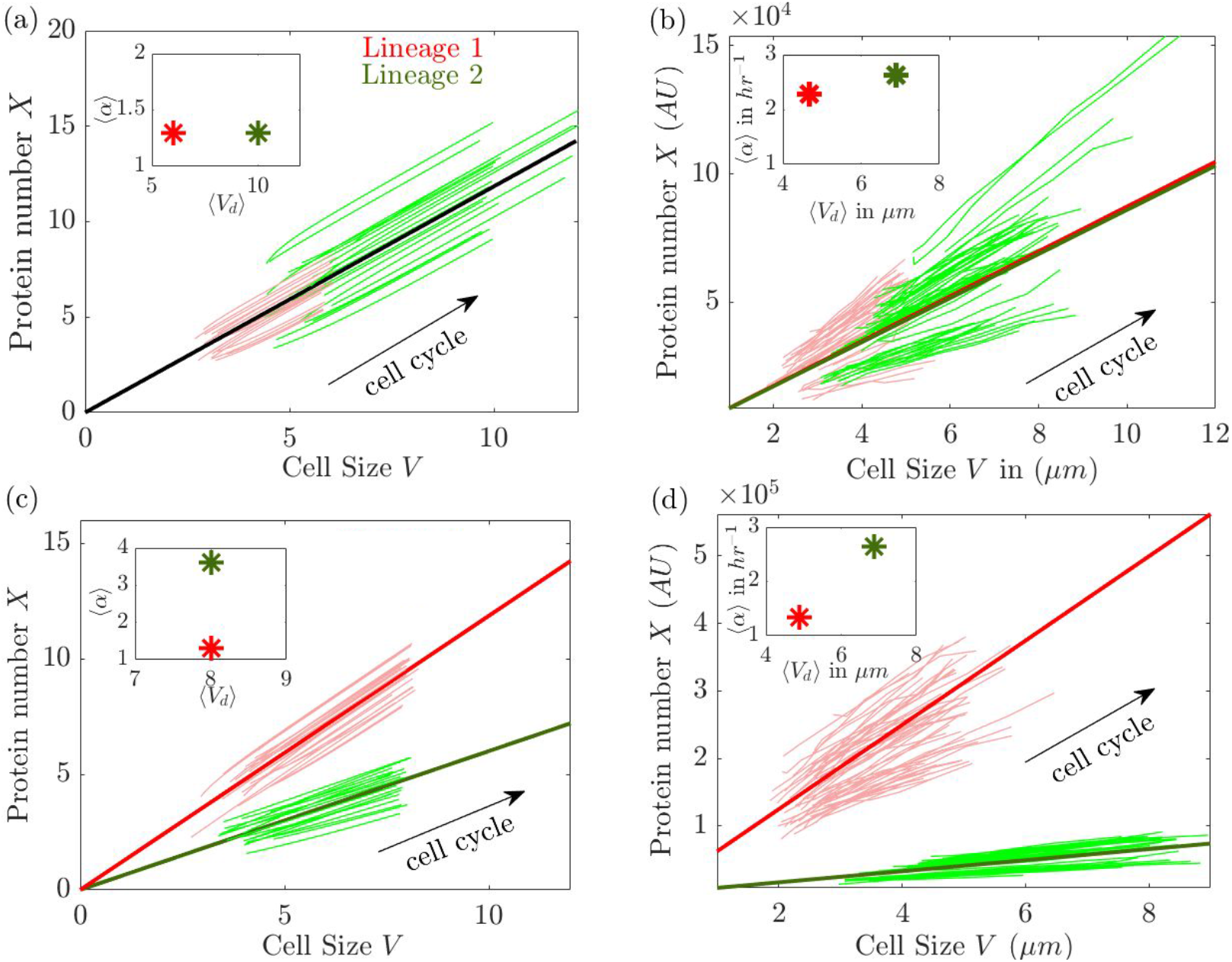
Degeneracy in exponential attractors: (a) & (c) Model; (b) & (d) Data. **(a)** Two lineages with different birth size (initial condition), obeying the same dynamics, maintain stable cycles of growth and division around different segments of the same attractor (dark green straight line, defining a fixed ratio of protein number to cell size). **Inset:** their average growth rate is the same, determined by their common attractor, but their average cell size differs according the region of the attractor. **(b)** Similar behavior of two lineages in experiment. **(c)** Two lineages obeying the same dynamical system but with different parameters, which give rise to distinct attractors (Dark green and red thick lines). Each attractor defines a different protein-to-size ratio, resulting in a different slope (Dark green and red thick lines). **Inset:** The two distinct attractors define distinct average growth rates, but because of degeneracy initial conditions can be chosen such that average cell size is the same. **(d)** Similar behavior of two lineages in experiment. Experimental data are on wild-type MG1655 *E. coli* at 30^°^ *C* in LB medium (protein expressed from *λ* − *pR* promoter) [15].

A comparable effect is observed in experimental data, as shown in Fig. 3(b). Here, two lineages measured within one experiment are presented in the same form as the simulation. The insets in Fig. 3(a) and (b) highlight that the two lineages share a similar growth rate—suggesting the same attractor—yet maintain homeostasis around significantly different average cell sizes, corresponding to different regions on the same attractor.

The example above illustrates degeneracy in the absolute values of cellular components, where trajectories within a single dynamical system—with fixed parameters—can originate from different initial conditions and sustain long-term homeostasis around distinct regions of the same attractor. However, a second layer of degeneracy arises when the system’s parameters themselves are modulated by internal perturbations, such as protein modifications, DNA configuration, and epigenetic or genetic regulatory modulation, as well as by external factors like temperature, chemical micro-environment, or pH. Each distinct set of parameters gives rise to a separate stable attractor with similar qualitative properties.

For example, in microfluidic devices, the growth dynamics of cells in different channels may follow distinct attractors due to subtle environmental variations. Even within the same micro-environment, epigenetic variability can shift a cell’s trajectory to a slightly different attractor. Since each attractor maintains fixed component ratios, projections onto two-component spaces appear as lines with different slopes. Figure 3(c) illustrates this effect in simulations, where two distinct attractors (black lines) support separate multi-generational trajectories (thin colored lines). The inset highlights that, in this case, the mean exponential growth rate differs between the two lineages, despite similar average cell sizes. This behavior is mirrored in lineages from one experiment, as shown in Fig. 3(d), where two measured lineages exhibit parallel dynamics within the same experiment.

### Homeostasis of the biological function in high-dimensional parameters space

The results and interpretation presented above suggest that cell size, absolute protein content, and even the precise exponential growth rate are not key regulated variables determining the cell’s functionality. Rather, these are “sloppy” variables that can exhibit persistent variability without compromising the biological function of growth and division homeostasis. This raises the question: what is the fundamental property that is maintained tightly regulated? Recent work suggests that homeostasis primarily preserves the stability of the growth and division process rather than individual variables. This stability is captured by the total average change in cell size over multiple cycles [30]. Specifically, the relevant compound variable encoding this stability is 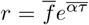, where 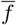 is the average division fraction, 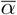 the average growth rate, and 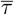 the average division time, all averaged over generations within a lineage. Although each of these averages varies across lineages, their co-variation induces compensation and can ensure homeostasis of growth and division.

To test the generaliuty of this principle, we analyzed a large dataset of single-cell measurements, averaging each lineage’s growth and division parameters over time and plotting the homeostatic variable *r* in the parameter space 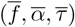. Figure 4 demonstrates how the data adhere to a well-defined manifold while displaying significant scatter over its surface. Each point represents a single lineage, with different colors denoting different experiments with distinct conditions. All data points adhere to the manifold, but are scattered along the individual coordinates - both within and across experiments. Very likely, this scatter reflects both different attractors (i.e., distinct sets of model parameters) and degeneracy within a single attractor.

**Figure 4.**
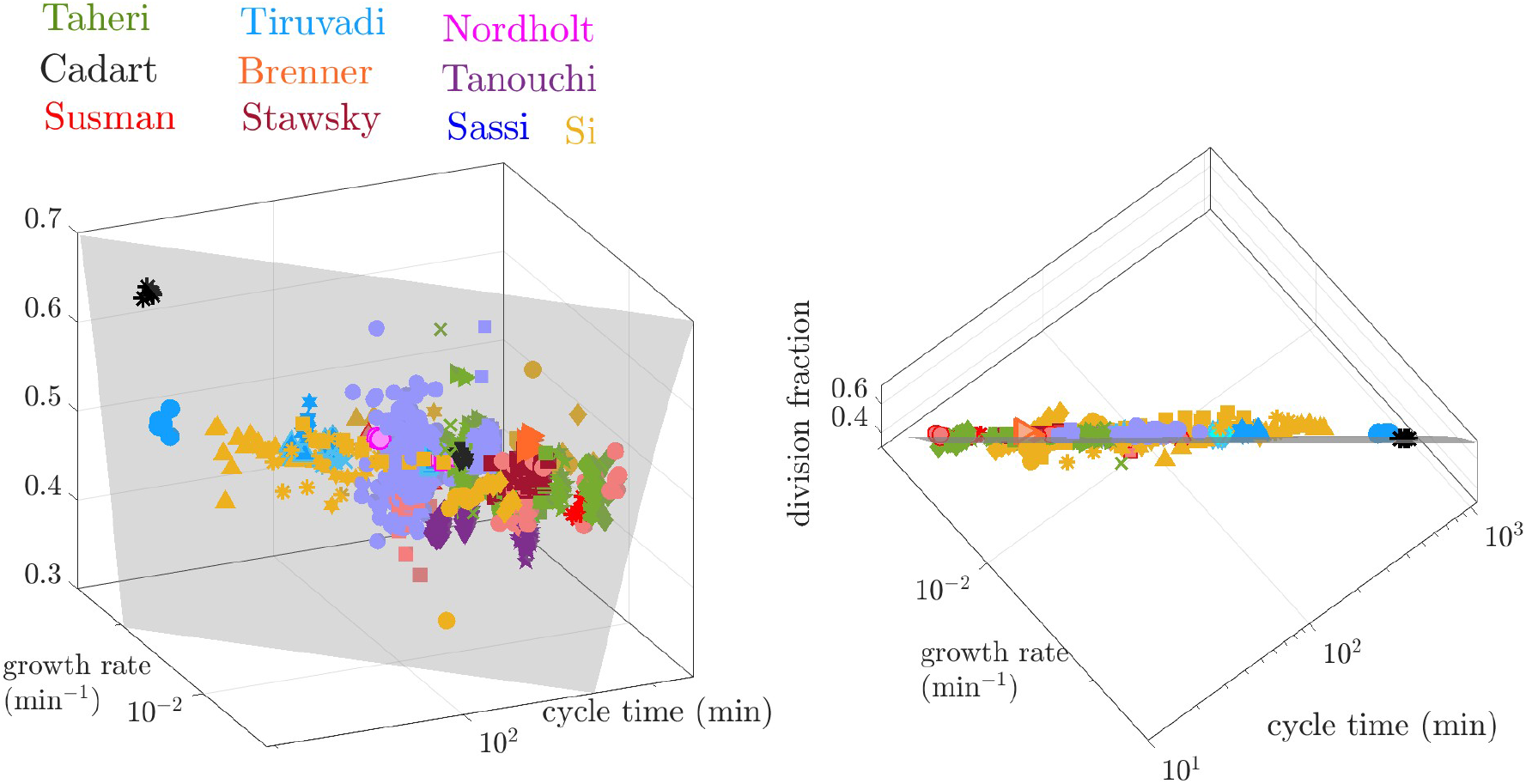
Plot lineage average of cell cycle time, growth rate, and division fraction from several experiments. Different colors correspond to different experimental groups as mentioned in the legend. Different shapes of the same color correspond to different experimental conditions, e.g., different media, different temperatures, etc. The data were collected from various experiments [15, 30, 20, 12, 14, 22, 23, 19, 21, 29, 31] with details provided in the Appendix.

Note that all the data presented in Figure 4 is *time averaged*; therefore type-1 variability is eliminated, and the scatter represents exclusively type-2 variability. The findings presented in this figure strongly support a hierarchical picture, where individual growth parameters exhibit high type-2 variability across lineages, and homeostasis operates at a higher functional level—preserving the stability of the growth-division process as a whole.

## Discussion

The complexity of biological systems gives rise to emergent phenomena, where system-level behavior cannot be fully understood or predicted by examining individual components separately. Accordingly, many fields in the life sciences have undergone a paradigm shift, driven by experimental advances that enable the simultaneous measurement of multiple variables. Notable examples include the ability to record the activity of hundreds to thousands of neurons over extended periods and to measure the expression of multiple genes within single cells. As these techniques continue to evolve, the structure of high-dimensional dynamics are being unraveled, providing new insights into system-wide phenomena. However, a fundamental challenge remains: how does high-dimensional microscopic *dynamics* give rise to the organized behavior of a system? [33, 34].

Real-time, simultaneous measurement of multiple variables (order 10) in single cells—while tracking growth and division—is rapidly advancing. Recent studies measuring both cell size and protein content have provided valuable insights into projections of the underlying high-dimensional dynamics. Understanding these dynamics, along with their attractors and stability, is crucial for deciphering homeostasis in complex systems, such as the cell, where multiple components interact.

Here, we presented a dynamic perspective of cellular homeostasis developed by examining directly the high dimensional space of multiple interacting components, and supported it by experimental data. The theoretical framework is based on a recent identification of a broad class of dynamical systems, in which stable exponential attractors with balanced biosynthesis emerge asymptotically, leading to the coordinated exponential growth of all cellular components [26, 27]. In this class, the attractor emerges from the dynamics without fine tuning of the interactions or the number of components. The cell is approximated as a growing, structure-less “bag of chemicals”, and the class includes all types of mass-action kinetics. It is therefore expected to be highly relevant to bacterial cells; indeed it is supported by a large set of data on different bacterial species in various conditions [28]. In mammalian cells, it may be necessary to take into account structure and segregation inside the cell. Systematic deviations from balanced biosynthesis were studied in proliferating human cells, and were found to be related to subcellular localization and organelle structure [35]. Nevertheless, we have included some eukaryotic cell data in our experimental tests as well, which follow similar behavior to bacterial cells. An extensive comparison between bacterial and mammalian single-cell data within the current perspective is a topic for future work.

Studying the properties of the dynamic attractor allows to identify two distinct types of phenotypic variability with different relationships to homeostasis. First, temporal fluctuations, which can accumulate and endanger long-term homeostasis: these are constrained by the phase space flow, which draws the trajectories back to the stable attractor.

Importantly, the attractor is a manifold defined by relationships among multiple variables - a “curve of balanced growth” [27]. Imperfect cell division plays a key role as a disruption that perturbs the smooth coordinated propagation on the attractor, a perturbation orthogonal to the curve, that crucially affects the statistical properties measured in single-cell experiments [28]. Following the disruption caused by division events, all variables are simultaneously drawn back over the next cycle. This effect suggests that the dynamic perspective obviates the search for dedicated regulatory circuits that separately detect deviations in specific variables, compare them to set points, and actively correct them. Instead, it highlights global properties which inherently stabilize a large number of system components simultaneously.

The second type of variability we identify is persistent individuality, which arises from properties of the attractor. In previous work, ergodic theory was applied to the projected space of concentrations, where the attractor becomes a fixed point [26]. However, in the physical space of absolute quantities, the dynamic attractor supports multiple layers of biological degeneracy. First, for a fixed set of model parameters, the emergent (unique) attractor has a flat direction that allows for many solutions with varying properties, such as different average cell sizes, while maintaining stable component ratios and concentrations. Second, for small changes in cell physiology or local environment that would result in slight variation of model parameters, neighboring attractors emerge, each supporting the high-level function of stable growth and division but with distinct physiological parameters, such as average growth rate. Note that the models we draw intuition from are coarse-grained, and include effective parameters that depends on hidden variables. This increases the chance of observing parameter variability under nominally uniform experimental conditions.

These intrinsic degeneracies introduce persistent variability that distinguishes individual lineages without disrupting homeostasis. Unlike temporal fluctuations, such variability does not accumulate over time. On the contrary, it may even support homeostasis by allowing the system to settle into different attractor regions where stability is maintained. Similar forms of degeneracy have been discussed in other biological contexts [36, 33, 9]. Cells can leverage this degeneracy to preserve key functional ratios and concentrations that ensure proper replicative homeostasis, while allowing variability in absolute quantities like protein content or cell size. Such degrees of freedom can enhance cellular adaptability to various environments and conditions, by defining “null directions” along which the system can easily drift while maintaining functionality, as has been observed in high dimensional neural dynamics during motor learning [37]. Our recent work has identified that, among different phenotypic variables, some exhibit larger persistent variability than others; this could correspond to a “sloppy system” structure [7], defining a hierarchy of significance to function, where the least significant variable are allowed to vary among individual cells while co-variation among them defines the more functionally relevant directions [30]. Understanding these analogies is topic for future theoretical and experimental work.

While our perspective emphasizes emergent homeostasis, we do not propose a strict dichotomy between control theory and dynamical systems theory. Control systems are, in fact, a subset of dynamical systems, and the two frameworks are deeply interconnected. For instance, feedback loops regulating cell size or protein content can be interpreted as coarse-grained approximations of the underlying high-dimensional system. Alternatively, a simple control circuit might represent a low-dimensional projection of a more complex dynamical system. It could also represent a separate module that is weakly connected to others [38]. A hallmark of complex biological systems is that they can be understood from multiple complementary perspectives [39]. Our goal is not to dismiss control-theory approaches, but rather to complement them with a dynamical systems view of homeostasis, where stability emerges from the system’s inherent structure rather than from explicit control mechanisms.

